# Macrophage-to-Myofibroblast Transition (MMT) – an adverse response to polypropylene mesh implanted for pelvic organ prolapse repair surgery in a non-human primate model

**DOI:** 10.1101/2025.11.03.686251

**Authors:** Marrisa A. Therriault, Katrina Knight, Srividya Kottapalli, Temitope Obisesan, Malini Harinath, Bryan N Brown, Pamela A. Moalli

## Abstract

Surgical repair of pelvic organ prolapse (POP) is often augmented by polypropylene mesh to provide mechanical support to the vagina and improve anatomical outcomes as compared to native tissue repair. However, POP repair surgeries utilizing PPM have complications (most often pain or mesh exposure into the vagina) in over 10% of cases. Previous work has demonstrated that tensioning of meshes with certain geometries (diamond and hexagon pores), results in both planar (pore collapse) and nonplanar (wrinkles) deformations, significantly altering textile properties and impacting the host response. To further investigate the impact of mesh deformation on the host response, we implanted mesh in a validated non-human primate model via sacrocolpopexy with stable flat (square pores, N=20) versus deformed geometries (mesh loaded on the diamond prior to implantation resulting in collapsed pores and wrinkles, N=20). To investigate the impact of tension independent of deformation, we implanted on and off tension (10N, N=10 in each group). We hypothesized that more stable geometries trigger a healing response that achieves homeostasis while deformed mesh, by increasing the amount of material in contact with the host, triggers a maladaptive remodeling response with the formation of myofibroblasts. After twelve weeks, we found that mesh deformations and the absence of tension increase the amount of mesh per area on the vagina (mesh burden) and reproduced clinical complications (mesh exposure and vaginal thinning). Interestingly, MMT cells, or myofibroblasts co-expressing a macrophage marker (CD68), were seen to significantly increase in response to mesh burden, as well as respond hyper-locally to the mesh fiber interface. We observed decreased collagen density and more immature matrix deposited in conditions with higher MMT cell presence, showing more disorganization in deposited matrix with increased mesh burden, and the loss of tension. TGF-β1, in both active and latent forms, increased with increasing mesh burden, and highest expression was observed in conditions precipitating the highest percentage of MMT cells, a possible mechanism of transdifferentiation. This study showed the importance of PPM mesh properties on mesh burden following tensioning, impact on MMT transdifferentiation, and the downstream effect of these changes on the host response and healing outcomes.

## Introduction

Pelvic organ prolapse (POP), or the descent of pelvic organs into the vagina, affects nearly half of women over 50 [1]. By age 80, 12.6% of women will undergo surgery to repair POP with a current annual economic burden of $1.5 billion [2, 3]. The demand for POP repair surgery is expected to increase by ∼50% by 2050 [4]. Minimally invasive sacrocolpopexy is the only current option for mesh augmented corrective surgery for those experiencing POP [5, 6]. This procedure requires a steep learning curve and can result in mesh-related complications, such as mesh exposure into the vagina and/or pain in ∼10% of cases [7]. Mesh complications have been associated with maladaptive remodeling response related to mesh deformation with mesh exposure due to stress shielding and pain related to fibrosis/encapsulation. It is thought that each response triggers a specific host response that, as of yet, remains incompletely understood.

Polypropylene meshes (PPM) used in POP repair surgeries are manufactured with textile properties (weight, porosity, pore size, and knit pattern) that impact the downstream foreign body response ranging from minimal to heightened inflammatory response with fibrosis, and encapsulation [8-12]. Notably, since the material is non-absorbable all responses are chronic. While most meshes are manufactured with pore diameters > 1mm and are intended to be implanted flat, properties often change *in vivo*. Indeed, when mesh is tensioned to restore anatomy, profound changes in mesh geometry occur including pore collapse, and wrinkles that alter properties by increasing material stiffness and material burden[13]. Our research has shown that diamond and hexagon shaped pores are particularly unstable [14] while square shaped pores maintain their geometry. When pores collapse to less than 1mm, it is problematic as the inflammatory response to neighboring fibers overlaps resulting in poor tissue in-growth, and fibrotic encapsulation [15, 16]. Clinically, meshes removed from women with complications that were initially implanted flat with open pores, upon excision, appear highly deformed with collapsed pores and wrinkles, and have minimal tissue integration [17-19]. Tensile testing of diamond and hexagon meshes under conditions mimicking sacrocolpopexy showed that most initial pore diameters >1mm completely collapsed with a complete loss of porosity at 10N – a load that is considered physiologic for surgical and *in vivo* loading conditions. The loss of porosity precipitates a heightened foreign body response [20]. Studies from our laboratory and others have shown that the immune cells localized to the implantation site of polypropylene mesh are predominantly macrophages that are phenotypically pro-inflammatory [17, 21, 22]. A maladaptive remodeling response ensues with collagen and elastin degradation and increased MMP activity leading to overall structural decline and mesh complications like exposure into the vagina and pain [11, 17, 23].

Fibroblasts, resident cells of the extracellular matrix (ECM), and their surrounding matrix exist in a reciprocal relationship, wherein the cells respond to biochemical and mechanical cues in the matrix that influence differentiation, migration, and ECM remodeling [24-27]. Previous research suggests that vaginal fibroblasts are highly susceptible to disturbances in the local mechanical environment, influenced by mesh burden [26, 28]. Fibroblasts become myofibroblasts hallmarked by the acquisition of contractile alpha-smooth muscle actin (α-SMA) fibers and matrix deposition in environments that are abnormally stiff or under supraphysiological tension like those imposed by mesh implantation [29, 30]. The high activity of matrix deposition and contraction by αSMA stress fibers characteristic of myofibroblasts contribute significantly to the on-going remodeling seen clinically as tissue contraction and fibrotic tissue deposition (the mechanism underlying mesh induced pain) [27]. TGF-β is released by macrophages in a pro-fibrotic (M2) phenotype, as well as by fibroblasts, even after the dampening of inflammatory stimulus [29, 31-33]. As shown by our lab and others, manipulation of the tension of an implanted polypropylene mesh results in an overwhelming myofibroblast response [21, 26, 28]. Previous studies have demonstrated macrophage signaling to fibroblasts can lead to their activation and differentiation to myofibroblasts [34-36]. Other studies have shown the existence of myeloid markers on myofibroblast cells, suggesting macrophage-to-myofibroblast transdifferentiation (MMT), and evidencing macrophage deposition of ECM [8, 23, 37, 38]. The dynamic and heterogenous nature of both cell types has been shown in response to implanted polypropylene mesh, however the exact presence of MMT cells and downstream outcomes of this communication and/or potential trans-differentiation on resulting healing outcomes is not clear and will be the focus of the current study.

We hypothesize that the primary driver of an optimal vs suboptimal outcome to an implanted PPM is related to its deformability upon tensioning with more stable geometries triggering a healing response that achieves homeostasis while deformed mesh by virtue of increasing the amount of material in contact with the host and increased stiffness, triggers a maladaptive remodeling response with the formation of myofibroblasts via MMT pathways. To date, however, studies investigating the host response to mesh have compared PPM products from different manufacturers, each with unique textile properties and mechanical properties, precluding the ability to isolate out impacts of mesh deformations [17, 22, 39]. To address this, response, we used a square pored lightweight mesh (Restorelle, Coloplast) generally associated with good outcomes and rotated it to the diamond configuration such that deformations (collapsed pores and wrinkles) would occur upon implantation in a POP repair model mimicking the impact of diamond pored products. Using this technique in previous studies, we effectively reproduced the complications commonly observed in women [21, 26]. To test our hypothesis, we aimed to define and compare MMT presence in response to a lightweight mesh (Restorelle, Coloplast) implanted flat versus deformed configuration in a primate POP repair model (sacrocolpopexy) [26]. Myofibroblasts respond to increased stress in a tissue, that can be the result of an increased amount of a stiff material like PPM vs that induced by placing the mesh and hence, the tissue on tension. To separate the impact of tension vs material burden, we compared the host response in the presence and absence of tensioning the mesh to the sacrum. For our outcomes we quantified macrophage and myofibroblast density, MMT cells and their localization relative to the mesh. To assess impact on downstream outcomes, we measured collagen content, GAGs, and matrix metalloproteinases (MMPs). We measured our endpoints at 12-weeks following POP mesh implantation as a reasonable timepoint in which tissue incorporation into the device can be expected to have occurred and the long-term inflammatory response established.

## Methods

### Animals

Rhesus macaque non-human primates (NHP) were used in a POP repair model as previously described. One animal served as an experimental unit. NHPs were housed in standard cages with 12-hour light/dark cycles and given water ad libitum with a scheduled primate diet supplemented with fresh produce. Demographic information was collected upon NHP arrival: age, weight, gravidity (number of pregnancies), and parity (number of vaginal births). For this study, retired breeders were used, allowing for the considerations of pregnancy, birthing, and age as contributing factors to POP representative of the clinical population. A pelvic organ prolapse quantification (POP-Q) score was obtained before, and 12-weeks after, surgery on each animal after sedation (ketamine hydrochloride: 10 mg/kg, xylazine hydrochloride: 0.5 mg/kg) as previously described^32^.

### Implantation strategy

NHPs were anesthetized (isoflurane 1-2.5%), intubated, and prepared for a sacrocolpopexy. Transection of level I (uterosacral ligament) and II (paravaginal attachments) vaginal support was performed prior to total hysterectomy in N = 48 animals (IACUC 16088646). Restorelle (Coloplast, Minneapolis, MN) was sutured to the anterior and posterior walls of the vagina in 2 predetermined geometries to define the impact of deformation independent of tension as described in Table 1: flat square pore mesh (R0) versus mesh loaded on the diamond, introducing pore collapse, and manually wrinkled (RD). These groups were loaded with (R0, N=10, and RD, N=10) and without (R0NT, N=9, and RDNT, N=9) tension. Meshes on tension were placed at 10N using a spring scale. Meshes without tension were not secured to the sacrum and left free following implantation on the vagina. N=8 of animals underwent the same surgery but no mesh was inserted (Sham group). Implantation groups were randomized on the day of the surgery and surgeons were blinded until the time of surgery. NHPs were anesthetized (ketamine hydrochloride: 10mg/kg, xylazine hydrochloride: 0.5 mg/kg) and received analgesics (buprenorphine hydrochloride: 0.03 mg/kg, meloxicam: 0.2mg/kg) before, during, and after surgery, per IACUC 16088646. Animals were monitored daily for signs of pain and distress and weighed bi-weekly by laboratory and veterinary staff. Vaginal mesh-tissue complexes were harvested at 12-weeks post-surgery, assessed for macro-scale measurements of complications (explantations and thinning). Then, full thickness mesh-tissue complexes from the anterior mid-vagina were divided for immunohistomorphology and biochemistry. Mesh tissue complexes from the posterior vagina were analyzed for mechanical properties in a separate study.

**Table 1.** Demographics from non-human primates used for implantations. Retired breeders were used for implantation in this experiment to represent the population of patients that undergo pelvic organ prolapse repair. Weight (kg) and age (years) showed no significant differences between groups. Parity, or viable pregnancies to vaginal birth was not different across groups expressed here as the * median and ^ interquartile range (25th-75th). POP-Q scoring, used clinically to measure the decent of the cervix from its original position on a scale of 1-4. Here, the POP-Q scores were not different between groups. *ANOVA reported. ^Interquartile range reported analyzed by Kruskal-Wallis.

|  | Weight (kg)* | Age (years)* | Parity^ | POP-Q stage^ |
| --- | --- | --- | --- | --- |
| **R0NT** | 11.1 ± 1.3 | 12.4 ± 3.2 | 3 (3-10) | 0 |
| **R0** | 9.4 ± 1.3 | 12.7 ± 1.9 | 4 (2-8) | 0 |
| **RDNT** | 10.0 ± 2.1 | 12.6 ± 2.2 | 5 (2-8) | 0 |
| **RD** | 9.1 ± 1.1 | 11.6 ± 2.5 | 3 (3-4) | 1 |
| **P-value** | 0.06 | 0.7 | 0.5 | 0.6 |

### Trichrome

To measure collagen density, formalin fixed and paraffin embedded anterior vaginal tissue sections (7um) were deparaffinized with xylenes followed by exposure to a graded series of ethanol (100-70%). and stained with standard Masson’s trichrome protocol. Sections were treated with Bouin’s solution and Weigert’s iron hematoxylin solution (to label nuclei) before Trichrome Stain AB Solution (Sigma Aldrich, Darmstadt, Germany). Tissue sections were imaged using a Motic Easy Scan Pro Digital Slide Scanner. Collagen density was quantified using open-source image analysis software QuPath [40]. The area of interest was defined by outlining the adventitia, the layer of vagina onto which the mesh is implanted and removing the mesh fiber area. Collagen density was defined by a threshold for blue intensity using a random forest pixel classifier. Collagen density was determined by diving the total blue pixels by the total area, providing a percent.

### Picrosirius Red

Fresh-frozen OCT embedded tissue sections (7um) were rinsed in DI H_2_O and stained in Picrosirius Red solution for 1 hour at room temperature with agitation. Picrosirius Red solution was prepared by mixing Direct Red 80 (365548, Sigma Aldrich, Darmstadt, Germany) with saturated aqueous picric acid (5860-16, Ricca Chemical, Arlington, TX, USA) to yield a concentration of 0.145mM. Tissues were decolorized in acetic acid water (1:2 glacial acetic water to DI H_2_O) for 10 seconds. Tissues were incubated once in 95% ethanol, twice in 100% ethanol, and thrice in xylenes for 1 minute each. Images were taken using Nikon ECLIPSE 90E upright microscope (Nikon USA, Melville, NY) under polarized light at 10X. Differential collagen deposition of thick versus thin fibers in vaginal adventitia was evaluated by calculating the amount of red vs green pixels using MATLAB.

### Hydroxyproline assay

Total collagen content was quantified from lyophilized and papain digested tissue samples with the hydroxyproline assay, using methods previously described[25]. It was assumed that hydroxyproline comprises 14% of the amino acid composition of collagen in calculations for total collagen content. The total collagen content was normalized to tissue dry weight.

### Glycosaminoglycan quantification (GAGs)

The amount of sulphated glycosaminoglycan (sGAG) in papain-digested vaginal tissues was quantified using the 1,9-dimethylmethylene blue assay as previously described [25, 26]. The amount of sGAG was normalized to tissue dry weight.

### Protein Quantification

Fresh-Frozen tissue was ground with mortar and pestle prior to being introduced to a high salt buffer with 1X Protease Inhibitor (04693132001, Roche, Basel, Switzerland) and 0.1% Triton-X. Samples were homogenized for 1 minute at medium speed using a TissueRuptor (QIAGEN, Hilden, Germany) and centrifuged for 20 minutes at 4C. Supernatant was collected and the tissue was homogenized with digestion buffer and placed at 4C and lightly agitated overnight. Samples were centrifuged for 20 minutes at 4°C and supernatant was combined with previous collected supernatant to make the protein isolate. Protein isolates were then analyzed using DC protein assay (Bio-Rad Laboratories, Hercules, CA) to empirically determine protein concentration.

### Collagenase activity

Collagenase activity was quantified from lyophilized protein extracts using a collagenase activity assay with fluorescein isothiocyanate (FITC)-labeled telopeptide-free soluble bovine type I collagen (Chondrex, Redmond, WA) as previously described[26]. Cleavage of 1ug of collagen per minute was recorded as 1 unit and collagenolytic activity was described as units/ml.

### TGF-beta quantification

Amount of TGF-β1 was quantified on protein isolates using a commercially available enzyme-linked immunosorbent assay (ELISA; R&D Systems, Minneapolis, MN) as previously described [26]. Total amount of latent TGF-β1 was approximated by subtracting the amount active TGF-β1 from total TGF-β1 content.

### Enzyme-linked Immunosorbent Assays (ELISAs) MMP Quantification

Protein isolates were calculated for each MMP assay (MMP1: 40ug, MMP2: 20ug, MMP8: 60ug, MMP9: 20ug, MMP13: 60ug). ELISA protocol was followed (R&D Systems, Minneapolis, MN) and samples were run in duplicate and MMP values were calculated from 4-PL standard curve. MMP quantity is reported as pg per ug of protein to normalize for the different amounts of protein necessary to run the assay.

### Meso-Scale Discovery U-Plex Assay (MSD)

Protein isolates were run on a commercially available custom multiplex electrochemiluminescence assay (Meso Scale Diagnostics; Rockville, MD) was used to determine concentrations of GM-CSF, MCP-1, MIP-1α, based on preliminary data identifying these factors as key players in the host response. Samples were run in duplicate with 80 µg protein.

### Immunofluorescent labeling

Fresh frozen tissues were quadruple labelled with CD68, vimentin, a-smooth muscle actin (αSMA), and DAPI. Full thickness anterior vagina was embedded in OCT prior to flash freezing on liquid nitrogen and sectioned (7μm) before storing at −80°C until use. Tissues were fixed in 4% paraformaldehyde at 4°C for 1 hour washed with agitation PBS 3 times for 5 minutes each. To reduce autofluorescence, tissues were incubated with combination of 10mM copper sulfate and 50 mM ammonium acetate at 37°C for 20 minutes. Tissues were washed and blocked with 4% donkey serum in 0.3% Triton X-100 in TBS for 30 minutes at room temperature. Primary antibodies of CD68 (ab955, monoclonal anti-mouse 1:50, Abcam), vimentin (NB300-223, polyclonal anti-chicken 1:100, Novus Biologicals, Centennial, CO), and αSMA (ab21027, polyclonal anti-goat 1:50, Abcam) were added to the tissue for 45 minutes at room temperature and then overnight at 4°C. Negative controls were concurrently incubated with blocking solution. Secondary antibody incubation followed with anti-mouse Alexa Fluor 647, anti-chicken Alexa Fluor 594, and anti-goat Alexa Fluor 647 each at 1:200 dilution for 2 hours at room temperature. Tissues were rinsed and washed in TBST followed by PBS. Slides were mounted with aqueous medium containing DAPI (Vectashield with DAPI, Vector Laboratories, Burlingame, CA). Whole tissue section immunofluorescent images were taken at 10X using a Nikon ECLIPSE 90E upright microscope (Nikon USA, Melville, NY). Images were analyzed using QuPath v0.4.3 to calculate the total number of positive labeling cells in the responding tissue omitting the area taken up by mesh fibers. To assess organization of the cellular response, we quantified percent of myofibroblasts per total number of responding cells at incrementally increasing concentric radius (50 um) from each mesh fiber.

### Statistics

Sample size was calculated based on previously published data assessing the myofibroblast response to deformed mesh in NHPs as a mechanism for poor remodeling outcomes. Based on the observed difference in myofibroblast counts between Sham and Predeformed groups in this study [26], a sample size of approximately 5 animals per group was determined to be sufficient to detect this difference with 80% power and α=0.05, using a two-tailed t-test. A sample size of 9 animals was sufficient to detect a difference in secondary outcomes (biochemistry and ELISAs-MMP/TGF**-**β). When normally distributed, samples were compared using a one-way ANOVA with a Bonferroni correction for multiple comparisons when appropriate. For non-parametric data or normally distributed with unequal variance between groups, Kruskal Wallis was used. Post-hoc comparisons were made if overall p<0.05. Significance values adjusted by Bonferroni correction for multiple tests when necessary. A two-way ANOVA was run on the cell distances from nearest mesh fiber to determine statistical significance between groups and distances. All statistical analysis was performed GraphPad Prism (Version 9.0.0 for MacOS, Boston, MA). The significance level was set to p<0.05.

## Results

### Gross morphologic assessment of excised mesh-vagina complex

Non-human primate demographics that are known risk factors for POP development were not significant between groups (weight p=0.06, age p=0.7, parity=0.5, POP-Q p=0.6, **Table 1**). Mesh-vagina complexes were excised at 12-weeks following implantation. Gross morphological assessment of measurable clinical outcomes was observed and recorded upon removal (**Figure 1**). Measurable clinical outcomes, reported in **Table 2** and depicted in **Figure 1**, indicate vaginal thinning (circled) - a potential precursor to a mesh exposure (arrows) through the epithelium. The location of these exposures was primarily at the apex, where mesh implanted on the anterior and posterior vagina converge. Vaginal thinning was a significant outcome seen in nearly 50% of the excised mesh-vagina complexes, more commonly in the RD and RDNT groups (p=0.07, **Table 2**), and adjacent to mesh exposures throughout the vagina. Exposures increased with increasing mesh burden (R0<R0NT<RD<RDNT, p=0.0492, **Table 2**).

**Table 2.**
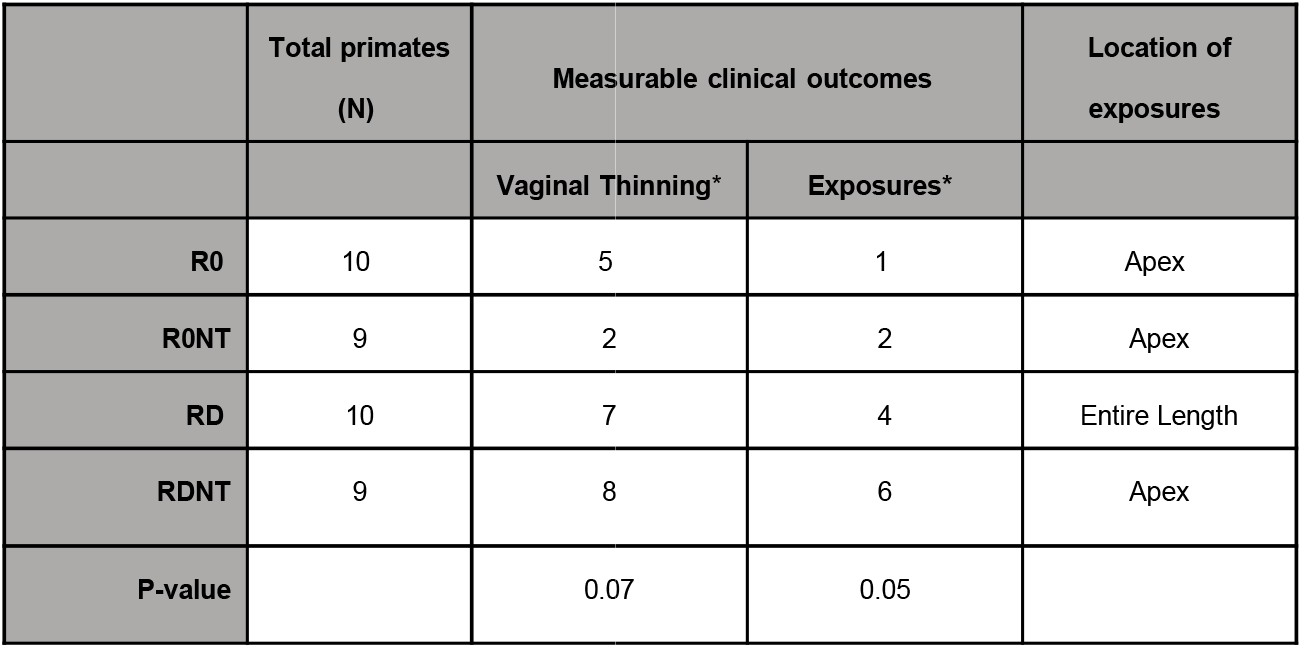
Quantification of clinical complications from increased mesh deformations at time of implantation in non-human primates shows an increase in complications. Total number of NHP used in this study is reported (N). Measurable clinical outcomes, indicated by vaginal thinning and exposure of the mesh through the vagina, is seen to increase with increased introduction of mesh deformations (R0<R0NT<RD<RDNT). The location of the exposures is reported, most of which being at the apex of the vagina, or where the mesh is secured to the vagina in sacrocolpopexy. *ANOVA reported.

**Figure 1.**
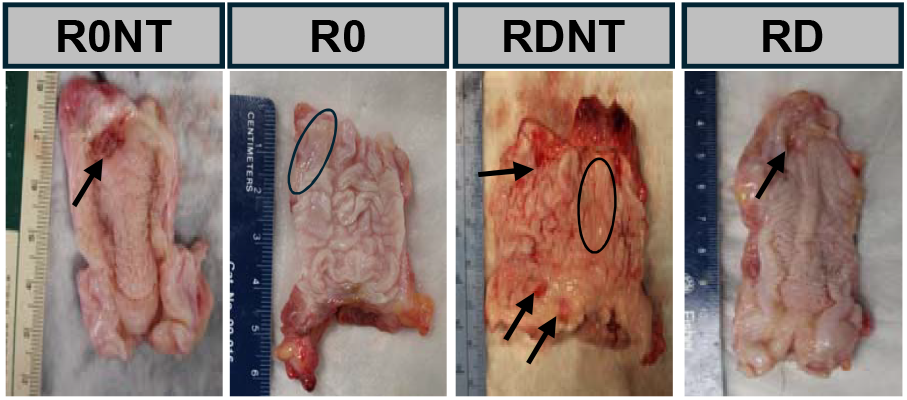
Planar (pore collapse) and non-planar (wrinkle) deformations intentionally introduced into a single light weight mesh resulted in clinically relevant complications. Sacrocolpopexy was performed on non-human primates with Restorelle implanted on the square in the presence (R0) and absence (R0NT) of 10N of tension, as well as mesh deformed with both planar and nonplanar deformations introduced upon implantation with (RD) and without (RDNT) 10 N of tension. Tension was measured with a spring scale prior to attachment of the mesh to the sacrum. Mesh tissue complexes removed after 12 weeks indicate vaginal thinning as indicated with circles and exposure of the mesh through the vagina indicated with arrows. Quantification of these clinical outcomes is seen in Table 2.

### Cellular response to PPM

The density of cells responding to the PPM in the adventitia showed no significant difference in cell infiltrate between groups (p=0.3, **Figure 2B**). Mesh area, determined as total area of adventitia occupied by mesh fibers, showed a significant difference between groups (p=0.0007, **Figure 2C**). R0 had the lowest mesh burden, followed by R0NT, RD, and lastly RDNT, emphasizing that the absence of tension further increases mesh burden. In particular, RDNT had significantly higher mesh burden than R0NT (p=0.02) and R0 (p=0.0004), but not RD (p=0.2).

**Figure 2.**
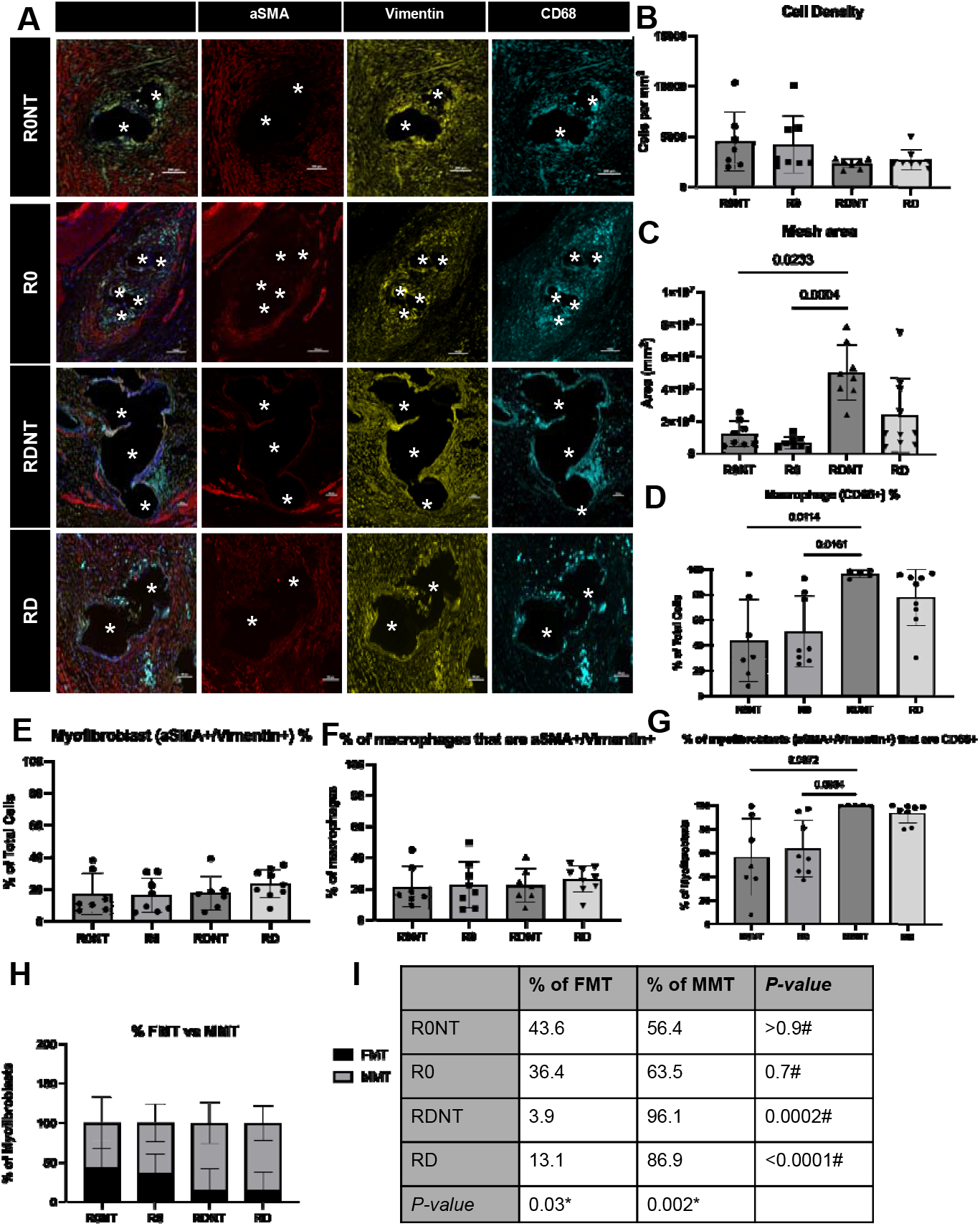
Myofibroblast and macrophage co-labelling indicates macrophage-to-myofibrob response to implanted mesh. A. Immunofluorescence labelling shows DAPI (blue), myofibrobla red), fibroblast marker (vimentin, yellow), macrophage marker (CD68, cyan). Imaged at 20X, Scal fibers indicated by astrix. B. Cell density, as measured by total cells in the adventitia (site of mesh not significant between groups (p>0.05). C. Mesh area, or area of the adventitia occupied by mes increased with introduced mesh deformations, thereby defining an increase in mesh burden with in deformation (p=0.0007). D. Macrophage percentage, or percent of responding cells labelling CD68 increased mesh burden (p=0.005). E. Myofibroblast percent, or percent of responding cells in the vimentin+aSMA+ was not statistically significant between deformation groups (p=0.4). F. % of ma with myofibroblast markers, or CD68+ cells that also labelled with aSMA+vimentin+, was not signi deformation groups (p=0.6). G. % of myofibroblasts that co-labelled with CD68, or aSMA+/Viment labelled with CD68+ significantly increased with increasing mesh burden (p=0.002). H and I. Perc myofibroblasts that co-labelled with a macrophage marker defined here as MMT (aSMA+Vimentin percentage of myofibroblasts that labelled with myofibroblast markers only were defined as FMT (The percentage of myofibroblasts that are MMT cells increased with deformation (p=0.0002) while FMT labelling myofibroblasts decreased with deformation (p=0.03). *ANOVA value reported. #Dun comparisons reported.

Macrophages were the primary cell type responding to the mesh in all groups. Macrophage (CD68+) percent of total cells responding increased with increasing deformation with R0 at 54%, R0NT at 48%, RD at 79% and RDNT 96% (p=0.003, **Figure 2D**). Significant differences were seen with R0NT<RDNT (p=0.01) and R0<RDNT (p=0.009).

Vimentin+αSMA+ are myofibroblasts that originate from fibroblasts and are classified as FMT, while the myofibroblasts arising from macrophages colocalize Vimentin+αSMA+ with CD68+ are classified as MMT. FMT cells as a percentage of total cells were not significantly different between groups (p=0.4, **Figure 2E**). MMT cells were also not significantly different across groups (p=0.6, **Figure 2F**) and comprised about 20% of total macrophages in all groups. However, as a percentage of myofibroblasts, MMT cells were seen to significantly increase with increasing deformation (R0NT<R0<RD<RDNT, p=0.002, **Figure 2G**). Out of all the responding myofibroblasts in each group, the percentage that co-labelled with the macrophage marker CD68 classifying them MMT cells was 56% in R0NT, 64% in R0, 96% in RDNT, 87% in RD. Significant differences were seen with R0NT<RDNT (p=0.007), and R0<RDNT (p=0.008) and no difference between RD and RDNT.

The total myofibroblast response was categorized by % of FMT or MMT cells responding, revealed a significant difference in the percent of myofibroblasts that are FMT (p=0.03) or MMT (p=0.002). Percent of myofibroblasts that were FMT significantly decreased with increasing deformation (R0NT 43.6%, R0 36.4%, RD 13.1%, RDNT 3.9%, p=0.03, **Figure 2I**), showing an increase in MMT cells with increased deformation (p=0.002, **Figure 2I**) Within each group, there was a significant difference in % of myofibroblasts that were FMT and MMT (p<0.0001).

### Organization of cellular response around mesh fiber

From the immunofluorescence labelled samples we were able to analyze the distance of a cell to the nearest mesh fiber. The percentage of each cell type at 100μm increments from the nearest mesh fiber (**Figure 3**) showed that, as expected, the macrophage response was highly localized to the mesh fibers (0-100um compared to all distance intervals >300um) in all groups (p<0.0001), with no difference between groups (p=0.01, **Figure 3A**). Myofibroblast were also measured by distance to nearest mesh fiber and were divided into FMT (αSMA+vimentin+) and MMT (αSMA+vimentin+CD68+). FMT myofibroblasts were found to be more localized to the mesh (0-100um) in the deformation groups (R0NT vs RDNT, p=0.01, R0 vs RDNT, p<0.0001, R0 vs RD, p<0.0001, **Figure 3B**). In the presence of deformation, MMT myofibroblasts were also found to be more localized to the mesh fiber (0-300μm) in R0NT, RD, and RDNT as compared to R0 (p<0.0001), with hyper-localization to the mesh (0-100μm) in R0NT (p=0.002), and RD (p=0.002) compared to R0 (**Figure 3C**).

**Figure 3.**
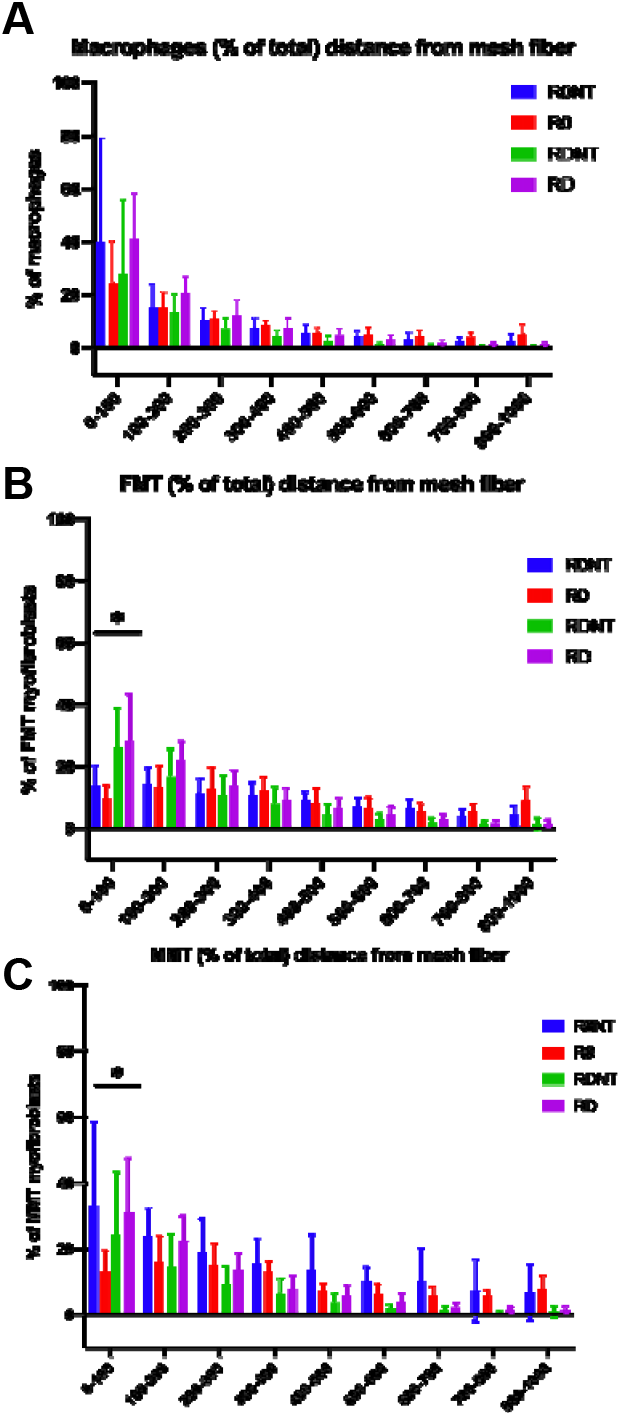
Cell distance from mesh fibers evidences a more localized cell response as measured by distance of cell to mesh fiber with increasing mesh burden. Immunolabelled samples for CD68, αSMA, and vimentin were analyzed for cell distance from the nearest mesh fiber. A. Macrophages (CD68+) were seen to have a robust response highly localized to the mesh fiber in all conditions (p<0.0001), with no significant difference between groups (p=0.1). B. FMT myofibroblasts (aSMA+vimentin+) were found to be more localized to the mesh with increasing burden (p<0.0001), with a more robust response seen closer to the mesh than in R0 (p<0.0001). C. MMT myofibroblasts (aSMA+vimentin+CD68+) were seen to be highly localized to the mesh fiber with increasing burden compared to R0 (p<0.0001) and observed closer to mesh fibers in increased deformation conditions than in the open pore conditions, despite tension (p<0.0001).

### ECM remodeling to PPM

Histologic quantification of collagen by Masson’s trichrome determined collagen density by quantifying the area occupied by collagen in the adventitia responding to the PPM (**Figure 4A and B**). There was no difference in collagen density between Sham and R0 - the mesh implanted flat with open pores (p>0.05). Collagen density was significantly different between groups (overall p<0.0001), observed as a decrease from Sham, in R0 (6.6%), R0NT (37%), RD (17.3%), and RDNT (57.2%, **Figure 4B**). Collagen density decreased significantly from Sham with the loss of tension (R0 vs R0NT, p=0.005) and independent of deformation (RD vs RDNT, p=0.0013, **Figure 4B**). Collagen deposition was quantified biochemically using hydroxyproline assay (**Figure 4C**). Collagen content significantly decreased from Sham across groups (R0=2.67%, R0NT 20%, RD= 25%, RDNT=19%, p=0.02), with a significant decrease in collagen observed in R0 compared to RD (p = 0.02, **Figure 4C**). GAGs content was quantified biochemically and expressed as percent difference from Sham showed significant differences (overall p<0.0001). GAG content increased from sham in groups without tension (R0NT 10%, RDNT 42%), but decreased in the groups with tension (R0 14% and RD 20%, **Figure 4D**). There was a significant increase in GAGs in RDNT compared to R0NT (p= 0.04) and RD compared to RDNT (p<0.0001, **Figure 4D**).

**Figure 4.**
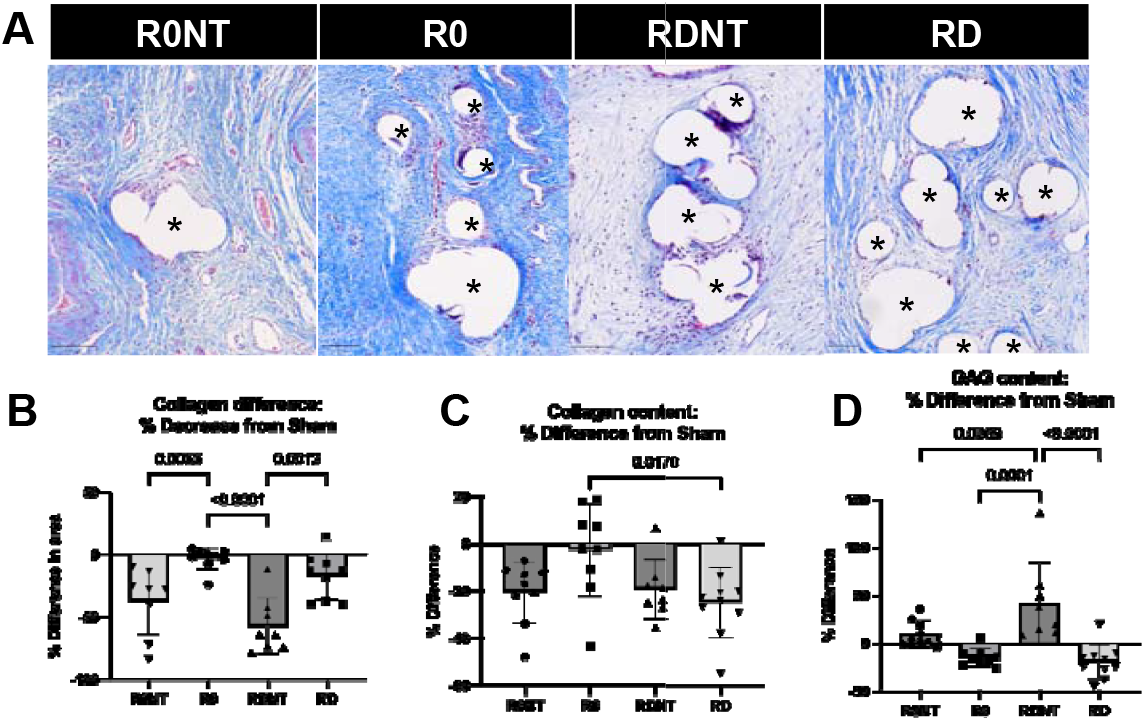
Collagen density increases in response to tension. A. Massons trichrome stained samples labelling collagen as blue, blood and muscle as pink, nuclei as purple, imaged at 20X. Mesh fibers identified by astrix. Scale bars 100um. B. Massons trichrome quantified as a percentage of adventitia stained blue (collagen) and shown as a % difference in collagen density from sham (control), showed a more significant decrease in collagen density in the no tension groups (p<0.0001). C. Collagen content determined biochemically by hydroxyproline assay, shown as a % difference from sham, found to be significantly different between groups (p=0.02). D. Glycosaminoglycan (GAGs) content quantified biochemically by GAG assay, shown as a % difference from sham, was found to increase in no tension groups, and decrease in tension groups (p<0.0001).

To further define characterize collagen in the groups, thin sections were stained with picrosirius red and imaged under polarized light. In this assay, thicker more mature collagen stain Red/Orange and less mature thinner collagen fibers stain yellow/green (**Figure 5A**). Percent of total collagen being mature, or red, was significantly different between groups (R0 97%, R0NT 95%, RD 99%, RDNT 96%, p=0.0004). There was significantly less mature collagen in the no tension group compared to the same group on tension (R0NT vs R0, p=0.02, RDNT vs RD, p=0.02, **Figure 5B**). An inverse relationship was seen for green, less mature fibers, with significant differences observed between the groups (R0 3%, R0NT 5%, RD 1%, RDNT 3%, p=0.0004), with no tension groups having a higher percentage of more immature newly deposited fibers than the tension group (R0NT vs R0, p=0.02, RDNT vs RD, p=0.02, **Figure 5C**). The ratio of mature (red/orange) to newly deposited/less mature (yellow/green) fibers showed the groups on tension (R0 and RDNT) having a higher amount of red/orange fibers compared to the groups without tension (R0NT and RDNT, p=0.0065, **Figure 5D**).

**Figure 5.**
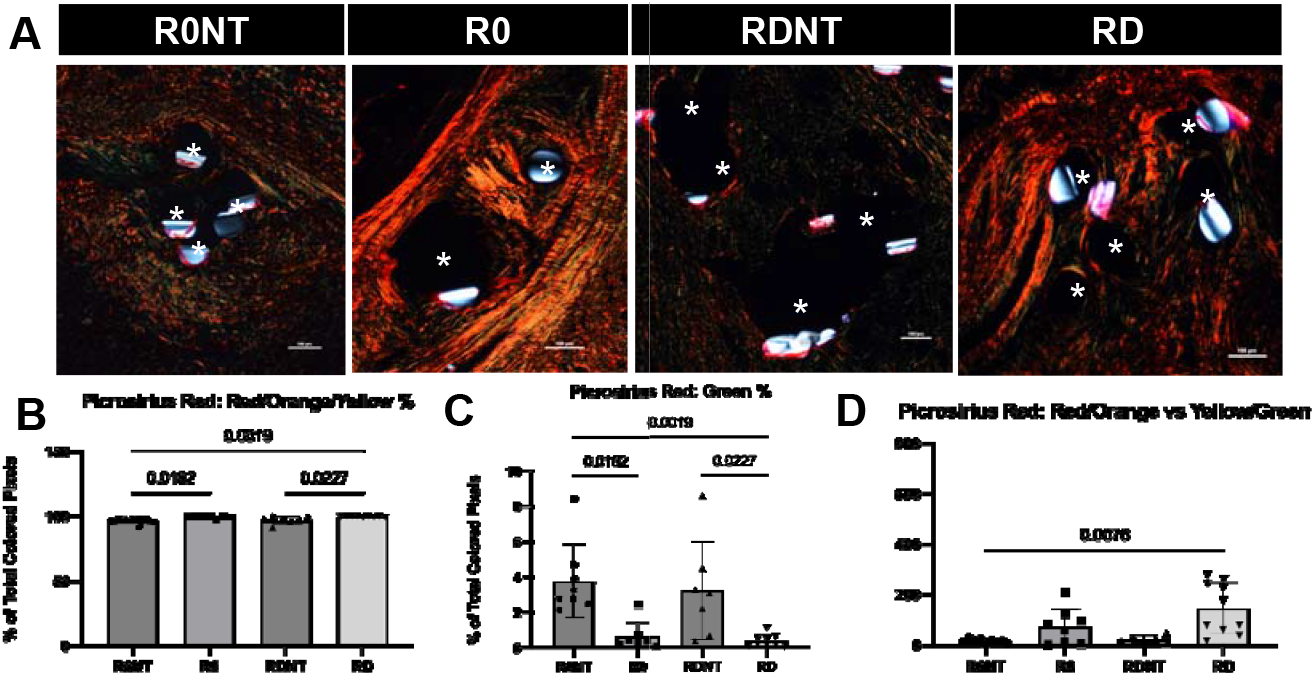
Tension induces increased collagen alignment and maturation. A. Picrosirius red stained and imaged using polarized light at 10X visualizes mature/thicker collagen as red, and less mature/thinner collagen as green. Scale bar 100um. Mesh fibers identified by astrix. B. Picrosirius red quantified using a previously published MatLab code defined the percentage of adventitia occupied by colored pixels, identified a significant difference in red or mature collagen deposition across groups (p=0.0004). With higher red/orange % in groups on tension. C. Green or less mature fibers were found to have a higher green % in the groups without tension (p=0.0004). D. Ratio of red/orange/yellow fibers vs green fibers was found to be significantly different between groups, with significantly more mature remodeling happening in the tension groups.

Matrix-degrading metalloproteases (MMPs) were quantified using enzyme-linked immunosorbent assay (ELISAs) and compared as a concentration (pg/mL) per μg of protein. Two types of MMPS - collagenases (MMP 1) and gelatinases (MMPs 2 and 9) were measured using ELISA to detect total MMP in mesh-tissue complexes. MMP-1 was found to be different across the groups (p=0.02), with an increase in MMP-1 in RD compared to Sham (p=0.02, **Figure 6A**). There was no difference in MMP-2 activity among the groups (p=0.18, **Figure 6B**). While there was a trend for a decrease in MMP-2 expression with a loss of tension, it did not achieve statistical significance (p>0.05). MMP-9, however, was significantly different between groups (p=0.002, **Figure 6C**). MMP-9 was increased in RDNT compared to R0NT (p=0.006), R0 (p=0.008), RD (p=0.001), and Sham (p<0.0001) consistent with the loss of matrix in this highest mesh burden group.

**Figure 6.**
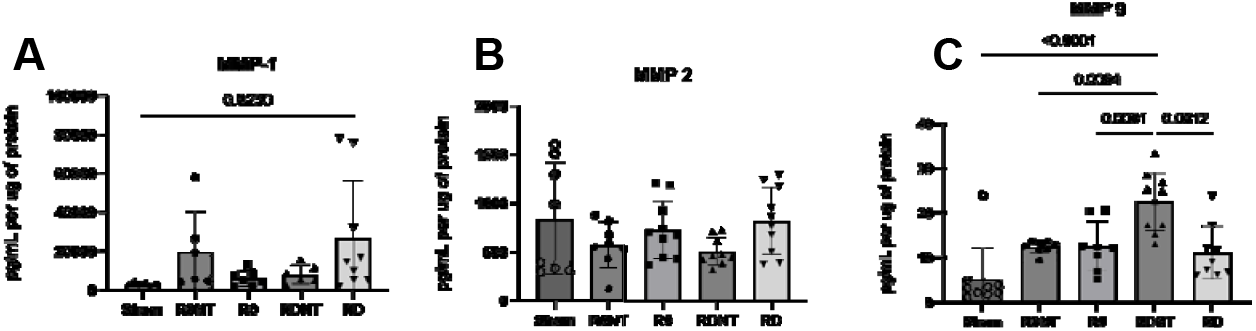
Matrix metalloprotease (MMP) -1,-2, and -9 levels in response to varied mesh conditions. MMPS-1,2,9 was quantified using ELISAs. MMPS A. MMP-1 was increased in all groups compared to Sham (p=0.02). B. MMP-2 was not found to be significantly different between groups (p>0.05) but does seem to be affected by tension. C. MMP-9 significantly increased with increasing mesh burden (p=0.002).

Since transforming growth factor-beta one (TGF-β1) is a potent immune modulator of myofibroblast transformation, it was quantified in both latent and active forms using ELISA (**Figure 7A** & **B**). Latent, or matrix-bound non-active TGF-β was found to increase in groups (p<0.0001), with a significant increase observed in RDNT (p<0.0001), R0NT (p<0.0001), R0 (p<0.0001), and RD (p<0.0001) as compared to Sham (**Figure 7A)**. Active TGF-β1, a form available for signaling, was significantly different between groups. Highest active TGF-β1 expression was observed in no tension groups - RDNT, and R0NT as compared to sham (p=0.003, <0.001, respectively, **Figure 7B**). R0 had significantly higher expression of TGF-β1 than R0NT (p=0.04, **Figure 7B**), but no significant increase from sham (p>0.999).

**Figure 7.**
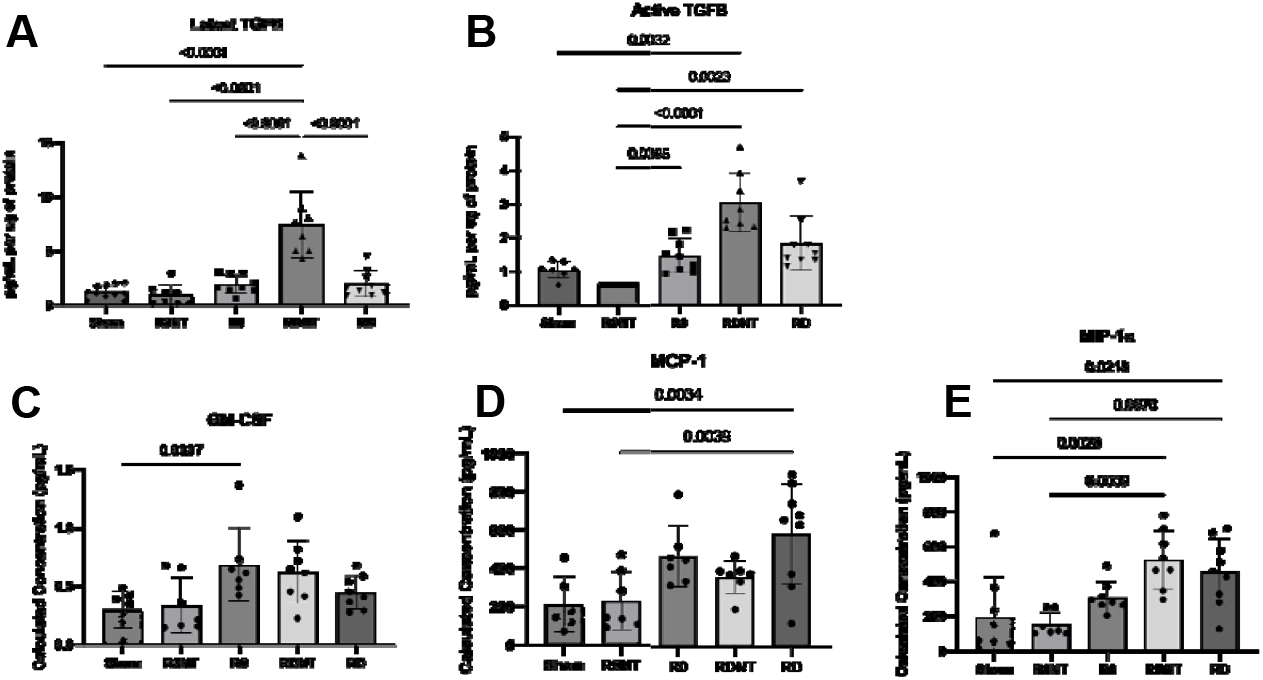
Macrophage-to-myofibroblast expressed cytokines increased with increased mesh burden. A. TGF-β1 expression, as measured by ELISA indicated expression of latent and active TGF-β1. Latent, or matrix-bound TGF-β1, increased with increased mesh burden (p<0.0001). B. Active TGF-β1 also increased with increasing mesh burden (p<0.0001). Expressed as pg/mL per ug of protein. C-E. Meso-scale discovery (MSD) assay revealed expression of macrophage chemokines like GM-CSF (C) MCP-1 (D) MIP-1a (E) expressed as calculated concentration as pg/mL. C. GM-CSF, or granulocyte-macrophage colony-stimulating factor, increased in all mesh groups compared to sham (p=0.01). D. MCP-1, or macrophage chemoattractant protein, increased with tension and deformation (p=0.001). E. MIP-1, or macrophage inflammatory protein 1, increased in all groups compared to Sham (p=0.02), and significantly increased with increasing mesh burden (p=0.0002).

Meso-scale discovery (MSD) assay was use to quantify macrophage secreted cytokines as a concentration (pg/mL) of protein isolates. Granulocyte-macrophage colony-stimulating factor (GM-CSF) is a cytokine that recruits and activates monocytes to the site of implantation. GM-CSF was higher in response to all mesh as compared to Sham with no difference between mesh groups (p=0.01, **Figure 7C**). Monocyte chemoattractant protein-1 (MCP-1) is expressed to attract and activate monocytes to the site of injury. MCP-1 was higher in all mesh groups relative to Sham and increased with tension and deformation (p=0.001, **Figure 7D**). Macrophage inflammatory protein-1 alpha is produced in macrophages as a mechanism of wound healing and is released by macrophages expressing a “pro-inflammatory” phenotype. MIP-1α was increased in the presence of mesh as compared to Sham (p=0.02) and increased significantly as mesh burden increased (p=0.0002, **Figure 7E**). Specifically, MIP-1α increased with deformation (R0NT vs RDNT, p=0.0009), but not with an increase in tension.

## Discussion

The goal of this study was to define the impact of mesh deformations in a clinically relevant non-human primate model to assess impact on the vaginal host response focusing on a pathological cell type, the myofibroblast. Previous studies have examined the effects of other mesh types (varying properties, manufacturers, etc), however, this study aimed to use a single mesh in two distinct geometries with and without tension. We showed that tension impacted mesh burden by reducing it, and mesh burden overwhelmingly drove the healing response to the mesh, and conditions with high mesh burden resulted in the most complications observed as vaginal thinning and mesh exposures, two clinically relevant complications following mesh implantation. The most important finding of the current study was that while macrophages, as expected, were increased with increasing mesh burden, myofibroblasts were also present with a surprisingly large percentage co-labelling with a macrophage marker. This signified a macrophage to myofibroblast transition (MMT) as opposed to a fibroblast to myofibroblast transition (FMT) was the predominant myofibroblast response with increased with mesh burden. MMT cells were highly localized to the mesh, where a macrophage traditionally responds. Measuring the healing response to the deformation groups we found that collagen density and maturity of deposited collagen was largely increased with tension, and decreased with deformation. Consistent with this finding, we observed higher expression of matrix metalloproteinases in the no tension, high deformation group (RDNT). Signaling of myofibroblast and macrophage recruitment cytokines evidenced an increase in expression of TGF-β1 and MIP-1α in the no tension, high deformation group (RDNT). Therefore, we found quite unexpectedly that tension has a mitigating effect on the host response, even in the presence of deformation. Moreso, we found that the highest percentage of MMTs were found in response to the highest amount of mesh burden and in conditions with complications. The predominant matrix-remodeling response was profibrotic in the presence of a mechanical stimulus (tension) and degradative in the absence of tension with high mesh burden and chemical signaling (TGF-β1) driving this response.

Previous work has been performed to define the textile properties of mesh that cause poor healing outcomes [41, 42]. These studies have associated high porosity, open pored, lightweight meshes with the most optimal healing response [15, 16, 22, 25]. However, for meshes that meet these criteria, the geometry of the pore must be taken into account as there is a risk for instability with loading *in vivo* that can alter properties and compromise outcomes. For example, highly porous lightweight meshes with diamond pores suffer a dramatic loss of porosity (<90%) with tensioning, suggesting these materials may not be suitable for urogynecologic repairs, where mechanical loading is required, like sacrocolpopexy [10, 26, 43]. Other work has defined the role of pore geometry and tension on responding cells [21, 26]; however, the current study is the first study to address the role of tension in the healing response as an independent factor from deformation.

We found that with each progressive deformation (R0<R0NT<RD<RDNT) there was increased mesh burden, as quantified by the amount of mesh in contact with the vagina. Using Restorelle, a lightweight, high-porosity mesh, R0 served as the “gold standard group” implanted flat, with open square pores, and tensioned. With each introduced deformation (R0NT: square pore, no tension; RD: introduced wrinkles, with tension, RDNT: introduced wrinkles, no tension) mesh burden increased. This increased burden was seen with an increase in clinically relevant complications, vaginal thinning, and mesh exposure through the vagina. Mesh exposures, a well-known mesh complication, occurred primarily at the apex of the vagina, where most of the load on the vagina occurs and hence, the mesh properties are most affected by deformation (R0 n=1, R0NT n=2, RD n=4, RDNT n=6). Additionally, the apex is the site where anterior and posterior straps of mesh meet (highest mesh burden) and the site of the hysterectomy incision which may heighten the host response.

The cellular response to increasing mesh burden illustrated a heightened macrophage response with increased deformation (RD and RDNT) consistent with our previous findings [21, 22]. However, when investigating myofibroblasts (all αSMA+vimentin+ cells) there was no significant difference between mesh implantation groups. Myofibroblasts are known to be highly proliferative and responsive to mechanical stimuli [29, 30]. In response to open pored flat mesh (R0), myofibroblasts formed a compact ring around each fiber. However, with increased mesh burden, this highly patterned response became disorganized suggesting there is less area to resolve around mesh fibers, and that mechanical signaling is now multi-directional due to increased mesh burden potentially causing the myofibroblasts to treat this as an immature wound and respond to the heightened degradation by macrophages.

In phenotyping the responding myofibroblasts, we found that a large percent of these cells co-labelled with CD68 (CD68+αSMA+vimentin+), a pan-macrophage marker, previously defined as macrophage-to-myofibroblast transition (MMT) cells. MMT cells have been defined in other organ systems in fibrotic environments, informing the capacity of macrophages to transdifferentiate to a myofibroblast phenotype and deposit fibrotic matrix as an element of the healing response [34-36]; however, their role in synthetic biomaterial soft tissue interactions is not clear. We observed the percentage of myofibroblasts that are MMT to be significantly higher than the percentage of the more traditional fibroblast-to-myofibroblast transition (FMT) cells with increasing mesh burden. MMT cells are seen highly localized to the mesh fiber, where macrophages typically respond, supporting their myeloid origin [8, 41]. This suggests that with increasing mesh burden, a higher percentage of myofibroblasts responding are MMT and therefore contribute to the maladaptive healing response.

Collagen density and maturity decreased progressively with mesh deformation, with the steepest decline occurring under conditions of lost tension. The accumulation of thin, disorganized, and immature collagen fibers suggests that the tissue is undergoing repeated cycles of degradation and repair rather than progressing toward resolution [44-46]. While MMP expression was elevated in mesh-implanted groups compared to Sham, the overall differences were modest, with the highest MMP-9 levels detected in RDNT, the group with the most disorganized collagen matrix. This pattern indicates that metalloproteinases, though contributory, may not overcome the dominant effect of persistent ECM deposition driven by myofibroblasts. Instead, the interplay of continuous paracrine signaling from macrophages and autocrine signaling from myofibroblasts appears to sustain disorganized matrix remodeling [38, 47].

In parallel, our data suggest that macrophage-to-myofibroblast transition (MMT) plays a central role in driving these maladaptive outcomes. A large fraction of myofibroblasts in high-burden conditions were MMT-derived, particularly in RDNT, where both cellularity was low and collagen remained immature. These findings align with the well-established role of TGF-β as a key mediator of MMT transdifferentiation, which we found to be most active in RDNT at 12 weeks post-implantation [34, 48]. Together, this points to a self-perpetuating loop in which macrophages, through MMT and sustained TGF-β signaling, contribute to both ECM deposition and disorganization. Whether these MMT cells phenotypically behave more like degradative macrophages or repair-oriented myofibroblasts remains an open question, but their presence at the mesh interface underscores their importance in the repeated injury–repair cycles characteristic of maladaptive remodeling.

Macrophage recruitment to the site of implantation was evidenced by increased expression of GM-CSF and MCP-1 in the mesh implanted groups compared to sham, however similar recruitment signaling between groups. Because this is a 12-week timepoint, macrophage recruitment is not expected to be ongoing, so this is indicative of a continual recruitment of macrophages longitudinally, and a shorter timepoint would further inform the increased macrophage presence in the groups with higher mesh burden. MIP-1α, or macrophage inflammatory protein, has been identified in wound healing environments released by macrophages to promote fibrotic tissue deposition [50]. Previous work has indicated that the surface topography of mesh may be altered by mechanochemical distress through tensioning and oxidation, and therefore this distress in vivo could play a role in the macrophage response to mesh [49-52]. Here, we found MIP-1α has a similar pattern to MMT cells, with increasing expression seen with increasing mesh burden, with the highest expression seen in RDNT. MIP-1α is released by macrophage and fibroblasts and could have a role in influencing the MMT transdifferentiation because both MMT cells and MIP-1α play a role in promoting matrix remodeling [44, 53].

In summary, our findings indicate that increased mesh burden, caused by mesh deformations, induces an overwhelming macrophage response that is hyper-local to mesh and induces cells that phenotypically express αSMA. In contrast to prior studies in which MMTs were profibrotic, we observed a novel phenotype characterized by a decrease in collagen density, increased matrix degradation, and deposition of immature matrix. This phenotype was most prevalent in the absence of tension indicating more of immune modulation potentially driven by TGF-β1 signaling - a known pathway for MMT transdifferentiation. A different response was observed when tension was applied with increased collagen organized in thicker/more mature fibers, and decreased GAG content, suggesting that mechanical stress signals a profibrotic pathway. Gross morphologic observations from this study showed that deformation and tension directly relate to mesh exposure through the vagina, a measurable clinical outcome for our study. Lastly, the spatial organization of the phenotypically non-classical macrophages (MMT) hyper-local to the mesh-tissue interface warrants future study of MMT cells and their role in the host response to implanted biomaterials. The mechanism by which MMT cells are activated in response to implanted biomaterials also warrants future study, and further investigation into the distinct features in the vagina as compared to other soft tissues.

## Limitations and Future Directions

Our study is limited in its scope, providing only a 12-week time point with no longitudinal information prior to or after 12-weeks. Though the non-human primate model is the most clinically relevant model, to further study the macrophage-to-myofibroblast transition a model with more frequent timepoints to provide more information on the recruitment, differentiation, and downstream outcomes of MMT cells responding to an implanted biomaterial. Continuing to identify the host response to PPM implanted for prolapse repair, we inform future devices and therapeutics to have the potential to improve healing outcomes clinically.

## Acknowledgements

The authors are grateful for the financial support provided by the National Institutes of Health R01 HD083383 grant funds to PAM. Research efforts for this publication were also supported by the National Center for Advancing Translational Science of the National Institutes of Health under award number TLTR001858 scholar funds to MAT. The content is solely the responsibility of the authors and does not necessarily represent the official views of the National Institutes of Health

## Data Availability

The data that support the findings of this study are available from the corresponding author P.A.M., upon request.

